# Busted: maternal modifiers of the triploid block involved in seed size control

**DOI:** 10.1101/2024.02.20.581014

**Authors:** Ruth Y. Akinmusola, Harvey Kilsby, Catherine-Axa Wilkins, Volkan Cevik, Rod Scott, James Doughty

## Abstract

The triploid block leads to seed abortion in crosses involving tetraploid Col-0 pollen. The genetic basis underlying this phenomenon is established in the endosperm and attributed to parental genomic imprinting. This research utilised the genetic variation in Arabidopsis to identify the genomic regions harbouring the maternal modifiers of the triploid block to produce viable large seeds. Distinct chromosomal regions were identified in Bla-1 and Tsu-0 accessions. The Bla-1 maternal modifier maps to the *TTG2* locus at the lower end of chromosome 2 to produce large viable seeds in response to a triploid block. Tsu-0 accession, on the other hand, recruits the *TTG1* locus on the upper arm of chromosome 5 as a maternal modifier of the triploid block. *TTG1* and *TTG2* mutations significantly increased the proportion of large viable seeds in interploidy crosses. Both genes are involved in transcriptional regulation in the flavonoid biosynthesis pathway. However, to regulate seed size in diploids, *TTG1* functions synergistically with auxin but does so independently of *TTG2*. This work contributed to the genetic framework for the *TTG1* and *TTG2* seed size roles.

**HIGHLIGHTS:**

- **Different Arabidopsis accessions recruit maternal modifiers to repress Col-killing in F_1_ triploids.**
- **These maternal modifiers may operate in the same pathway, such as the flavonoid biosynthesis pathway or other interconnected pathways such as auxin.**
- ***TTG1* and *TTG2* generally increase F_1_ triploid survival but in an accession-dependent manner.**
- ***TTG1* differentially exhibits a strong positive additive interaction with auxin to increase diploid seed size.**
- **The *TTG1*/*TTG2* roles in diploid seed size control appear to have diverged somewhere in the auxin branch of the flavonoid biosynthesis pathway.**

## INTRODUCTION

Angiosperm seeds originate from a double fertilisation event: one sperm cell fertilises the egg cell of the megagametophyte to produce a diploid (2n) embryo, and the second sperm cell fertilises the diploid central cell to form the triploid (3n) endosperm (*1*). The entire process of seed development in Arabidopsis consists of the morphogenesis and maturation phases. A mature seed contains the embryo, the genetic material of the next generation, and the endosperm, with both being protected by the seed coat. The coordinated growth of the endosperm and embryo in the maternal seed coat tissues has a profound impact on the final seed size (*2, 3*). Seed size is an important component of seed yield and is a key determinant of evolutionary fitness (*3*). Diverse signalling pathways controlling seed size in Arabidopsis and rice have been recently reviewed (*4*). Other factors such as hybridisation barriers and environmental cues can also affect seed size, but the latter plays a lesser role in seed size (*4, 5*). Hybridisation barriers limit gene flow both within and between different species, and this plays a major role in speciation. In terms of reproductive isolation, these barriers can act pre or post-zygotically. Postzygotic barriers only occur after fertilisation events resulting in offspring inviability and weakness, which can be seen as lethality or embryo abortion (*6*). Importantly, disruption of endosperm development is a major phenomenon underlying postzygotic hybridisation barriers in seed plants, and this occurs in both interspecific and unbalanced ploidy crosses (*7, 8*).

The genetic basis of the strong postzygotic isolation in a triploid block is established in the endosperm, the nutritive tissue nourishing the embryo during seed development. The endosperm (2m:1p) is dosage-sensitive, strictly requiring a maternal to paternal genome ratio of 2:1 to develop and function correctly due to the existence of genomic imprinting in the endosperm (*9*). The result of interploidy hybridisations deviating from the 2:1 is not only dependent on the direction of hybridisation but also on the parent accessions. Many Arabidopsis accessions (sometimes referred to as ecotypes in terms of their primary habitat) are tolerant when their diploid mothers are crossed with their respective autotetraploid, whereas only a minority of the accessions display paternal triploid block (Bolbol, 2010). Substituting Col-0 with the other accessions as seed parents results in a high level of variation in seed survival and lethality, indicating that Col-0 pollen is only aggressive when paired with some specific accessions. Endosperm overproliferation and extremely delayed cellularisation are the developmental aberrations underlying the Col-killer syndrome in the submissive accessions (*10*). However, the accessions resisting the Col-killer to produce a high percentage of viable seeds possess some maternal modifiers that can restrict the excessive proliferation of the endosperm during seed development (Bolbol, 2010). This maternal modifier is not a complex trait, it is dominant and mendelises, thus allowing for its analysis in the progeny (Bolbol, 2010); these characteristics indicate that the genetic candidates underlying maternal rescue in Arabidopsis can be mapped.

Different models explain the seed lethality or viable large seeds resulting from a triploid block. The parent conflict theory centres on the allocation of maternal and paternal resources to the offspring to drive parent-of-origin gene expression (*11*). The differential dosage hypothesis argues that the expression level of a transcript controlling viability or lethality in the triploid offspring is dosage-dependent, and the expression is largely influenced by parental contribution (*12*). These two models do not sufficiently explain the origin of the imprinted genes, but these could be from the gametophytes, sporophytes, endosperm, and the seed coat. Another model is consistent with interactions between the developing endosperm and seed coat in early seed development. This includes various pathways acting during the delay in endosperm cellularisation to suppress seed abortion. One of the critical pathways repressing seed lethality that results from paternal excess interploidy crosses is the flavonoid biosynthetic pathway (FBP), where mutations of a maternally expressed regulatory gene (*TTG2*) and an enzyme-encoding gene (*TT4*) bring about earlier endosperm cellularisation (*10, 13*). These common roles of these FBP mutants in maternal rescue in the triploid block have been attributed to auxin redistribution in the seed (*13*). However, their exact mechanism(s) of interaction with auxin in seed size control is not yet understood.

Here we report that *TTG1* also plays a maternal role in the triploid block. *TTG1* loss in response to Col4x pollen leads to increased seed survival in an accession-dependent manner. Though *TTG1* functions upstream of *TTG2*, both genes play different roles in seed size in diploids. This difference is demarcated by auxin signalling, suggesting that the two genes may regulate different auxin pathways in determining seed size.

## RESULTS

### Distinct genomic regions mediate maternal rescue in Arabidopsis accessions

The strength of maternal rescue against Col4x pollen in Arabidopsis is accession-dependent (Figure S1). Some accessions like Col-0, Cvi and RLD mothers are highly submissive to Col4x pollen resulting in a high percentage of shrivelled seeds with low mean seed weight. However, other accessions like Tsu-0, C24, Bla-1, Kas, Per and Ws resist the killing activity of Col4x to produce viable offspring, resulting in high mean seed weight. The genetic basis of this variation on the maternal side is poorly understood. Therefore, we sought to map the genomic regions containing the maternal rescue genes in Tsu-0 and Bla-1. These accessions have pre-existing recombinant inbred lines (RILs) with distinct Col-0 mosaic backgrounds for QTL mapping.

The composite interval mapping method of QTL analysis detected a major QTL on chromosome 5 that has a significant effect in controlling maternal rescue in Tsu-0 x Col-0 RILs (Figure 1A). This broad QTL overlays two SNP markers (c5_04011 and c5_08563) on the upper arm of chromosome 5. A separate QTL mapping in Bla-1 x Col-0 RILs detected two distinct QTLs underlying maternal rescue in the Bla-1 accession. The 8.06 Mb main effect QTL in the Bla-1 accession resides on the lower arm of chromosome 2 and is flanked by c2_09429 (9.43Mb) and c2_17606 SNP (17.61Mb) markers (Figure 1B). An epistatic QTL was also detected on chromosome one, but this smaller locus is only significant for seed death and is bounded by these SNP markers c1_20384 and c1_25698. These results suggest that the main effect maternal rescuer against Col4x in Bla-1 is located on the lower arm of chromosome 2 and may participate in an epistatic locus with another gene. The two QTLs possess additive effects, indicating that an increase in Tsu-0 or Bla-1 alleles in each respective region accounts for high mean seed weight in the RILs. In contrast, a substitution with Col-0 alleles increases seed lethality at both loci.

**Figure 1:**
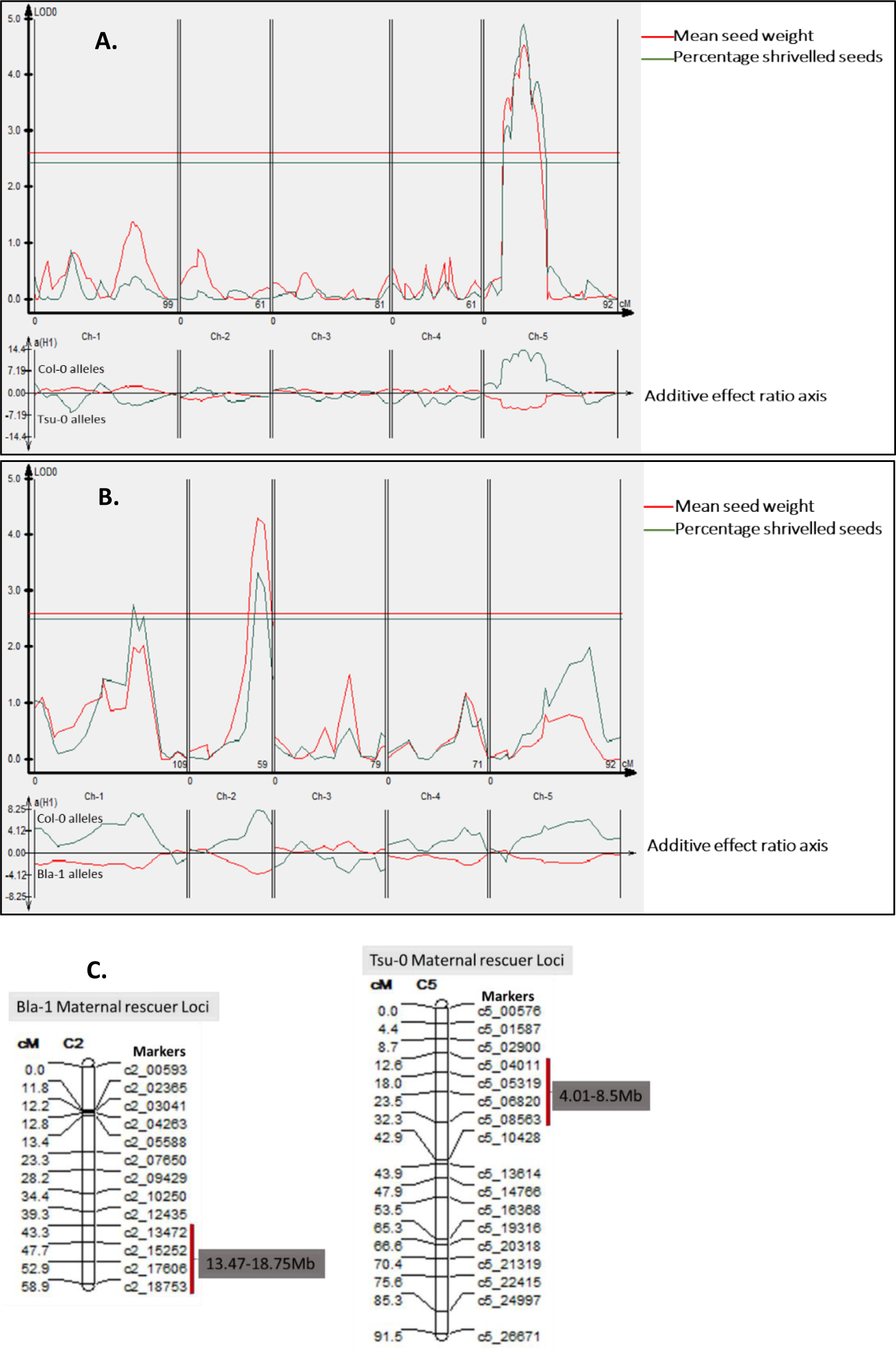
Two distinct QTLs underlie maternal rescue in the Bla-1 and Tsu-0 accessions. The upper graph shows the main QTL effects. Significant QTLs were detected above the LOD threshold score of 2.5 and an estimated *p* < 0.001 (the blue and red horizontal lines) derived from the y-axis. The corresponding *A. thaliana* genomic positions according to chromosome (Ch) number is shown in centiMorgans (cM) on the *x*-axis. For comparison, the smaller graph at the bottom is the QTL effects window, showing additive or dominant effects from Col-0 (positive y-axis) and other alleles (negative y-axis). Ch represents chromosome. Figure 1C: A chromosome Map showing the positions of the two QTLs.

The main effect QTL detected in BLa-1 is similar to *DSL1 (DR STRANGELOVE 1)* previously reported in the Ler-0 accession (*10*). The two distinct QTLs in Tsu-0 and Bla-1, indicate that different Arabidopsis accessions recruit various genetic factors to mitigate Col4x paternal excess. This within-species variation in genetic tolerance between diploids and tetraploids controls gene flow and could have arisen due to evolutionary divergence in gene expression affecting seed fitness or gene regulatory network (*12, 14, 15*).

### QTL refinement to narrow the Tsu-0 QTL locus

### Development of a secondary mapping population

The major QTL controlling maternal rescue in the Tsu-0 accession overlays two SNP markers (c5_04011 and c5_08563) on the upper arm of chromosome 5 (Figure 1). An analysis of the 4.55Mb sequence flanked by these two markers revealed that this QTL interval contains 1,197 genes. A set of indel markers polymorphic between Col-0 and Tsu-0 alleles were used to narrow down the QTL interval (Table S1). The Tsu-0 genome was aligned with the TAIR 10 Col-0 reference genome using the IGV software to hunt for the sequence deletions that occur in Tsu-0 reads within the 4.55Mb QTL region. These Tsu-0 deletions were filtered by their proximity to protein-coding genes, intactness, and size. Only deletions with a size between 100 bp to 2kb was retained, based on the expected product sizes between Col-0 and Tsu-0 (Table S1). These indel markers were used for genotyping in the subsequent analysis. Following the QTL location estimation and marker design, three RILs displaying extreme phenotypes for high mean seed weight were chosen to derive a secondary mapping population based on their recombination breakpoints within the QTL interval. Thus, these recombinants served as the donor parents in the development of some BC1F2 near-isogenic lines (NILs) through marker-assisted backcrosses.

### Narrowing down the 4.55Mb Tsu-0 QTL to a 0.48Kb interval via marker-assisted backcrosses

To estimate the physical position of the Tsu-0 maternal modifier gene within the 4.55Mb QTL on chromosome 5, a BC_1_F_1_ population was generated by introgressing the genomic region of a recombinant RIL into a diploid Col-0 background. These BC_1_F_1_ individuals exhibited the maternal rescuer trait, indicating that the Tsu-0 maternal modifier of Col-killing is dominant (Figure 2A and B). The BC_1_F_2_ individuals generated by selfing an individual BC_1_F_1_ segregated to show a high level of continuous phenotypic variation for mean seed weight (Figure 2A and B). Similar trends were observed in the BC_1_F_3_ individuals (Figure 2B). Each of the BC_1_F_2_ and BC_1_F_3_ populations had individuals with significant extreme rescuer (very high mean seed weight) and extreme submissive phenotypes with very low mean seed weight (Figure 2A and B).

**Figure 2:**
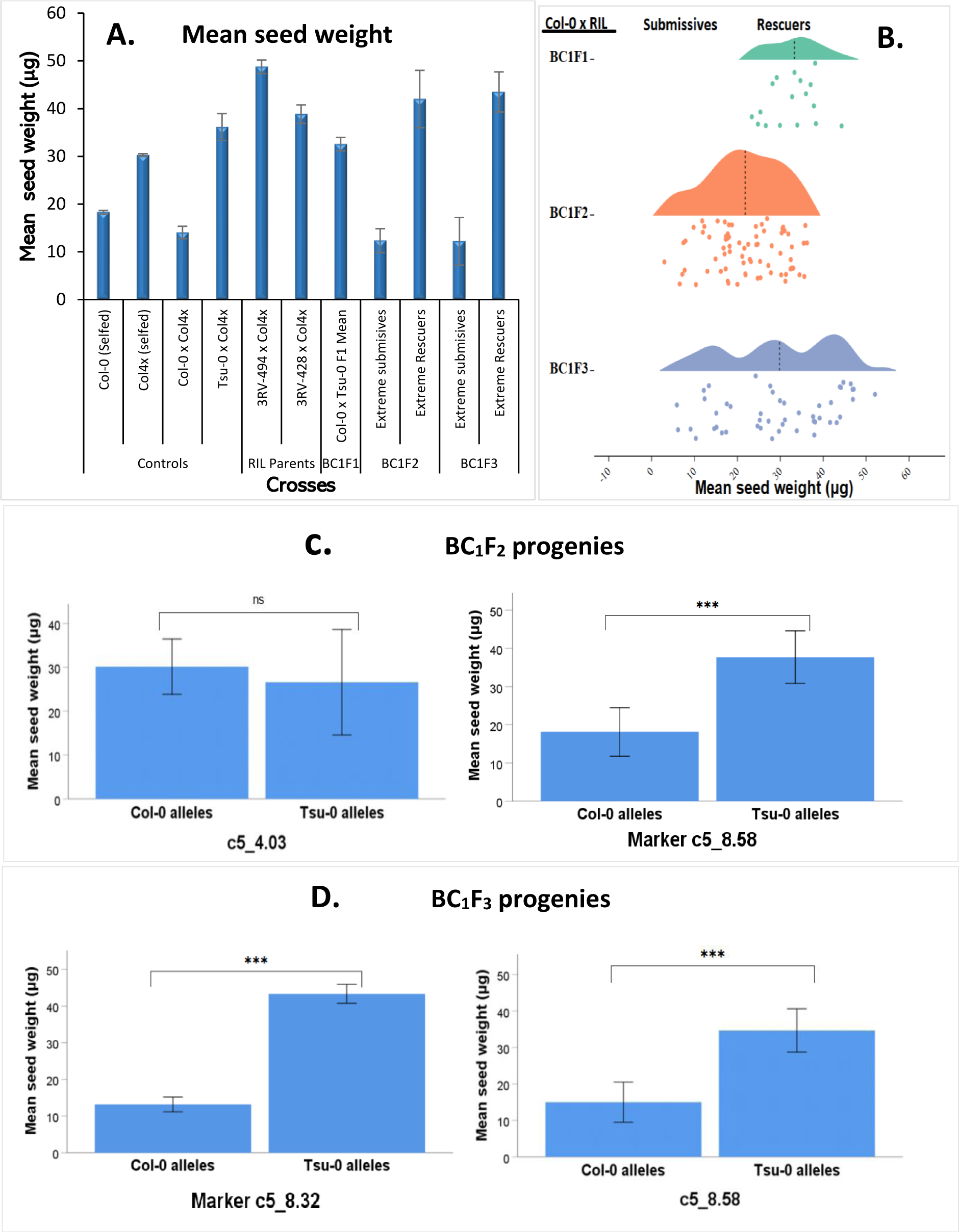
Two markers at the QTL peak (c5_8.32 and c5_8.58) are linked to the maternal rescue phenotype in the backcross progenies. A: The combined mean seed weight phenotype of the extreme rescuer and submissive BC_1_F_2_ individuals were also compared with the parental controls. Col-0 is the recurrent parent, and 3RV-428 is the donor RIL parent. B: General distribution of the maternal rescue phenotype (Col-0 x RIL) in the various backcross progenies (BC_1_F_1_, BC_1_F_2_ and BC_1_F_3_, respectively). D and E: Markers c5_8.32 and c5_8.58 are associated with increased mean seed weight phenotype in the progenies with Tsu-0 alleles. The error bars are the standard error of the mean. *: significant at p< 0.05; **: significant at p<0.01; ***: significant at p<0.001 and ns: non-significant.

The extreme individuals for both phenotypes were selected from the population to increase accuracy when validating their genotypes. A total of 16 individuals significantly distinguishable from the BC_1_F_2_ population were genotyped with nine indel markers. These markers have been previously designed to span less than 500Kb intervals within the QTL region and are based on deletions in Tsu-0 compared to Col-0, the reference genome (Table S1). The whole QTL interval is bounded by markers c5_4.03 at the beginning of the QTL and c5_8.58 at the QTL peak. The segregation ratios of the heterozygous alleles for each marker were excluded from the Chi-square analysis of marker segregation distortion (Table S4). This is because an equal level of homozygosity (AA=BB) is expected to occur in the parental alleles in a F_2_ population. In contrast to the other markers, c5_4.03 and c5_8.58 fit into the expected equal segregation ratio of homozygous Tsu-0 and Col-0 alleles at the 0.05 probability level (Table S4). However, only marker c5_8.58 at the QTL peak is associated with the maternal rescue phenotype based on the association of Tsu-0 alleles with a high mean seed weight and the association of Col-0 alleles with a low mean seed weight in the BC_1_F_2_ population (Figure 2C).

Furthermore, the segregation data of a BC_1_F_3_ population derived from selfing a recombinant BC_1_F_2_ individual was also used to estimate the chromosomal position of the Tsu-0 maternal modifier. A new indel marker, c5_8.32, was introduced between markers c5-8.20 and c5_8.58 to further delineate the region between these two markers in the BC_1_F_3_ population (Table S1, Table S4 and Figure 2D). Following genotyping, the genotype and phenotype data of the BC_1_F_3_ population was ranked according to the mean seed weight of the extreme progeny groups evaluated. Like the BC_1_F_2_ progenies, the markers used to screen the BC_1_F_3_ population displayed a segregation distortion towards one of the parental genotypes, starting from the beginning of the QTL (4.03Mb) to 8.20Mb (Table S4). However, two of these markers at the end of the QTL (c5_8.32 and c5_8.58) fit into the expected equal segregation ratio for Col-0 and Tsu-0 alleles in the BC_1_F_3_ population (Table S4). An increase in Tsu-0 alleles accounts for seed viability at these two markers, with c5_8.32 significantly associated with a higher mean seed weight, suggesting a tighter linkage to the candidate gene. The Col-0 alleles at this locus are associated with a significantly lower mean seed weight in the submissive progenies (Figure 2D). Thus, given these data, the Tsu-0 maternal modifier gene lies within the 0.48Mb interval between markers c5_8.20 and c5_8.58 (Tables S4C and D).

### The narrowed 0.48Mb Tsu-0 maternal modifier locus is enriched with genes involved in growth regulation

To screen the narrowed BC_1_F_3_ population identified in the previous section for potential candidate genes, all the genes in this narrowed QTL region were considered. The physical position of the indel markers at the extreme ends of this 0.48Mb region was used to retrieve all the candidate gene lists and gene models from Arabidopsis TAIR 10 genome via the UCSC genome browser. This retrieved gene list revealed 122 genes, and all these were analysed using the singular enrichment analysis tool of the agriGO portal (GO analysis toolkit and database for the agricultural community)(*16*). All the candidate genes were screened for significant associations between the traits they control and significant gene ontologies using overrepresented biological processes, cellular component, and molecular functions. The GO terms were parsed using their frequency, p values (<0.05), dispensability in a cluster, uniqueness, and their false discovery rate (FDR) values. Among all these criteria, p-values and FDR provided a clear-cut margin to identify statistically significant GO terms within each GO category (Table S2).

A summary of the GO analysis across all GO categories using the agriGO interface revealed 63 significant GO terms (<0.05) associated with different traits such as trichome morphogenesis and differentiation, hair cell differentiation, leaf morphogenesis and epidermis development. Highly significant GO terms with p values and FDRs lesser than 0.001 were used to screen all the GO terms analysed, using overrepresented biological processes, cellular component, and molecular functions. Biological processes constituted the largest bulk of the most significant GO terms (Table S2). Among these three terms, molecular function and cellular function seem to be the weakest in analysing these candidate genes as they are not directly linked to any trait (Table S2). The five most significant biological processes (BPs) topping the GO analysis are positive regulation of growth (GO:0045927), hair cell differentiation (GO:0035315), trichome differentiation (GO:0010026) and leaf morphogenesis (GO:0009965) were used as the final set of candidate genes. Each of these most significant BPs contains a relatively small number of candidate genes when compared to the other GO terms (Figure 3).

**Figure 3:**
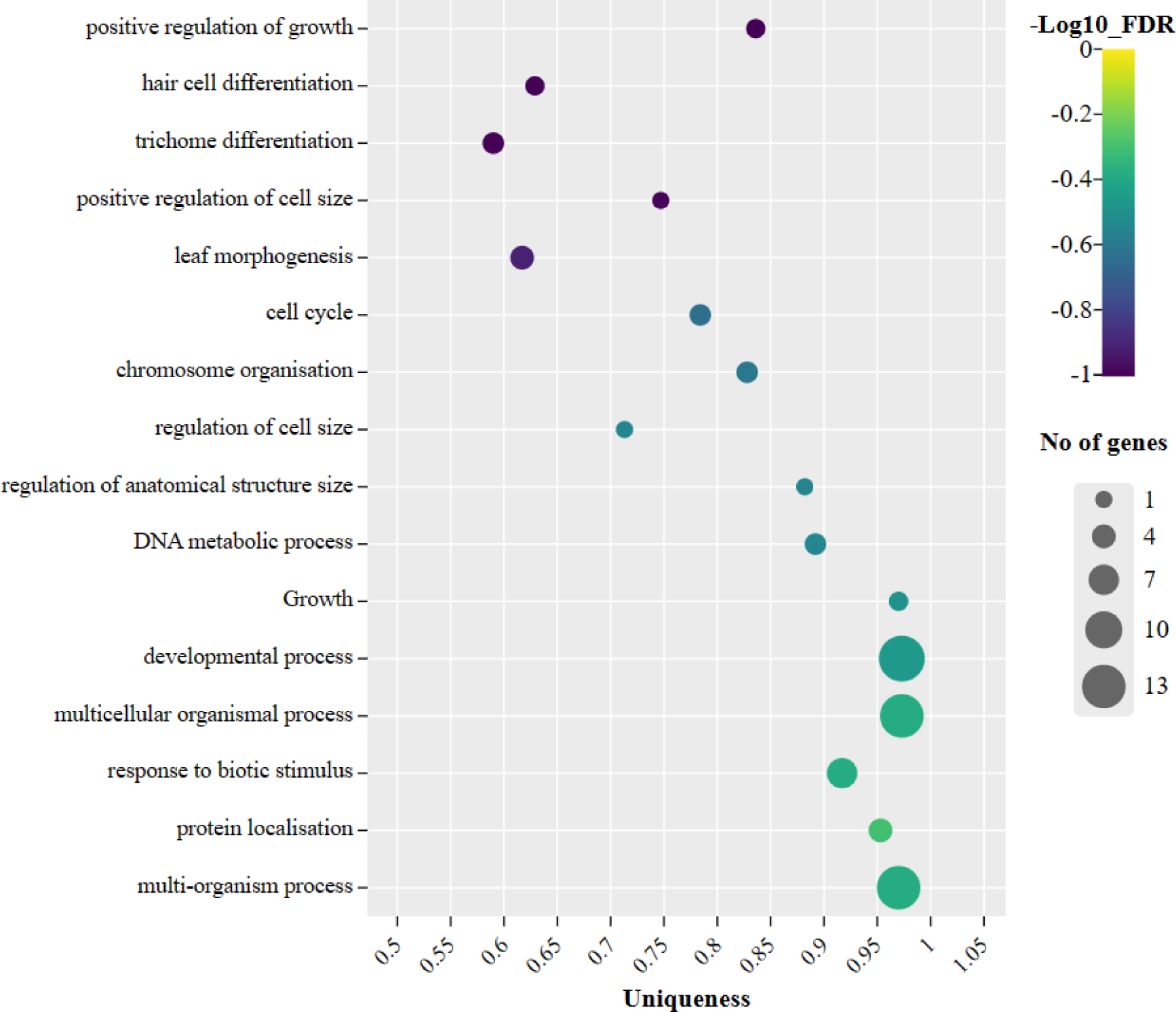
A relatively small number of genes are involved in the most significant BP terms within the narrowed QTL. The GO terms are clustered together in the plot. The Large bubbles represent many genes, whereas the small bubbles contain a small set of candidate genes. FDR represents false discovery rate.

### TTG1 is the major candidate gene at the 480 kb gene interval

The final list of genes in the narrowed QTL interval contains three genes in the final filtered gene set with a combined total of 10 variants (Table S3). These three genes are represented in the most significant GO terms in table S2. Among these, only *ATXR6* is fully expressed in the pollen and is involved in chromatin silencing. *ATXR6* encodes a H3K27 monomethyltransferase and its mutation results in plants with no physical defects, but they possess reduced H3K27me1 *in vivo* and the transcriptional activation of repressed heterochromatic elements (TAIR description). It interacts with the RNA processing factor (*RDR6*), a component of the RdDM to regulate transposon expression (*17*). However, mutation in the RdDM and its associated components only suppress paternal activity in F_1_ triploid killing (*18, 19*).

*BIN4* and *TTG1* share similar GO terms in biological processes such as trichome development (Table S3). *BIN4* encodes a plant-specific nuclear protein that is an integral component of the DNA topoisomerase VI complex in plants and is vital for cell endoreduplication and genome integrity in Arabidopsis (*20*). However, the functional loss of *BIN4* or any other part of the other plant topo isomerase VI components, results in the upregulation of the genes involved in DNA damage response (*21*). Thus, *BIN4* mutants usually display severe dwarfism with reduced cell and organ size and ploidy to about 25% of the size of the corresponding wildtype plants (TAIR description, (*21*). Based on these facts mentioned above, *BIN4* is unlikely to participate in the maternal control of chromatin modifications or DNA methylation in the triploid block.

*TTG1* encodes an evolutionarily conserved WD40 repeat protein in plants and its function is pleiotropic in the control of seed pigmentation, seed coat mucilage deposition, anthocyanin accumulation, trichome patterning, root hair patterning and even more recently the participation in flowering time regulation (*22-25*). *TTG1* is the transcriptional regulator for *TTG2*, a WRKY transcription factor that is involved in the maternal reduction of F_1_ lethality in Ler and Col-0 triploids (*10*). *TTG2* requires an intact *TTG1* to function in the flavonoid biosynthesis pathway and both genes share some similar phenotypes such as yellow seeds (*26-28*). Both genes are maternally expressed and *TTG1* was chosen as the final candidate gene based on its interactions with *TTG2*.

### *TTG1* mutation influences the survival of triploid F_1_s in an accession-dependent manner

Although Tsu-0 is heavily utilised in several mapping studies in arabidopsis (*29-35*), mutants in this accession are rare. This study generated a new *ttg1* CRISPR-Cas9 mutant in the Tsu-0 background using a *pHEE401E* vector with two guide RNAs. The *transparent testa* and *glabrous* phenotypes this novel mutant displays are consistent with the published panel of strong *TTG1* mutants (Figure 4). The maternal rescue potential of the *ttg1* mutant in Tsu-0 background was compared with a published panel of *ttg1* mutants in submissive Col-0 and Ler backgrounds (Figures 5 and 6, Table S5). Col4x pollen was chosen as the most potent pollen killer, whereas Tsu4x was selected as a mild pollen killer to assess the differences or changes in F_1_ triploid seed survival. *TTG2* mothers were also compared as controls. The maternal disruption of *TTG2* function has been reported to suppress the killing activity of Col4x pollen in F_1_ triploids (*10*).

**Figure 4:**
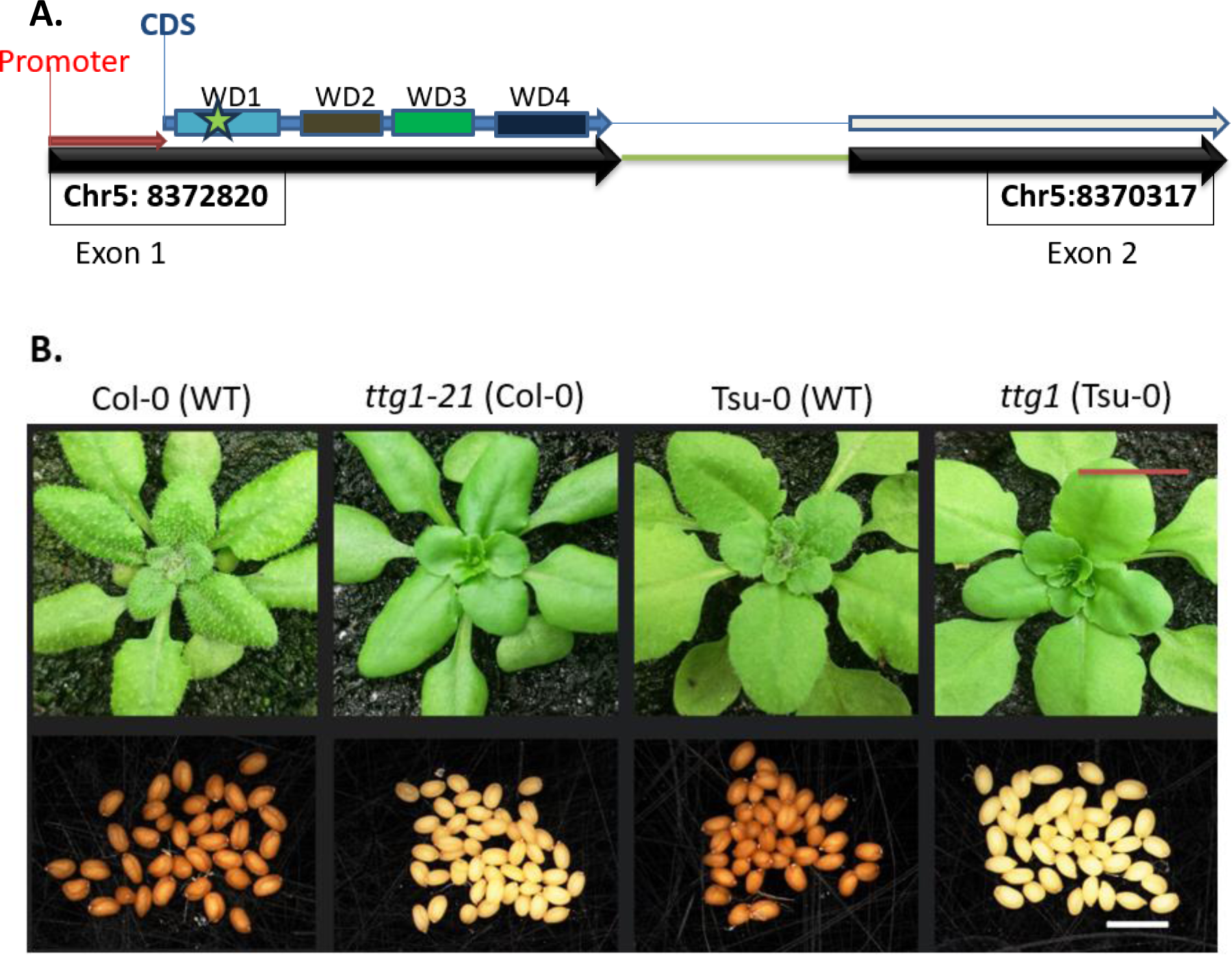
Phenotypes of the *ttg1*(Tsu-0) mutant. A: The corresponding genomic position of the *ttg1*(Tsu-0) mutation in base pairs (bp). The star represents the start location of the CRISPR-Cas9 mutation in Tsu-0 (Chr5: 8372730). The horizontal red arrow at the beginning indicates the 60bp *AtTTG1* promoter region (Chr5: 8372820-Chr5: 8372879). WD refers to the WD repeats in *TTG1,* which are four in number: WD1 (74-118), WD2 (130-170), WD3 (173-211), WD4 (262-302). The repeat features were extracted from UniProt. B: The ‘*glabrous*’ and ‘*transparent testa*’ phenotypes of the *ttg1* allele in Tsu-0 compared with the *TTG1* mutant in Col-0 background. The presence of a yellow and transparent seed coat depicts the *transparent testa* phenotype, while the absence of trichomes indicates the *glabrous* phenotype. Seedling images were taken three weeks after planting. The red scale bar is 1cm, and the white scale bar is 2mm.

**Figure 5:**
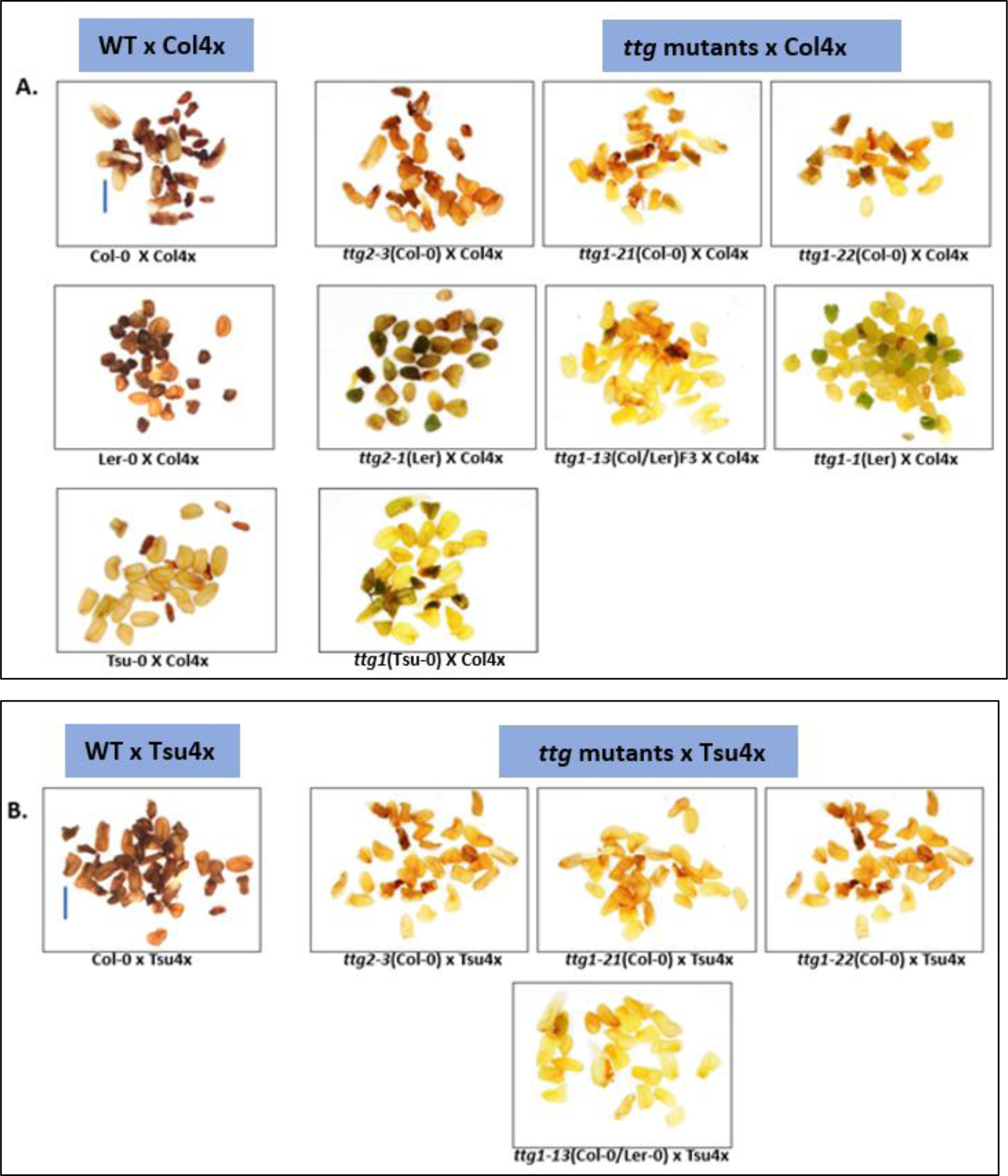
The maternal loss of *TTG1* results in an increase in plump seeds except in the Tsu-0 accession. The images are the triploid F_1_ seeds generated from *ttg1* mutants [2x X 4x] inter-crosses. The mother comes first in all crosses. All the images are in 20x magnification with a 1mm scale bar on Col-0 x Col4x and Col-0 x Tsu4x.

**Figure 6:**
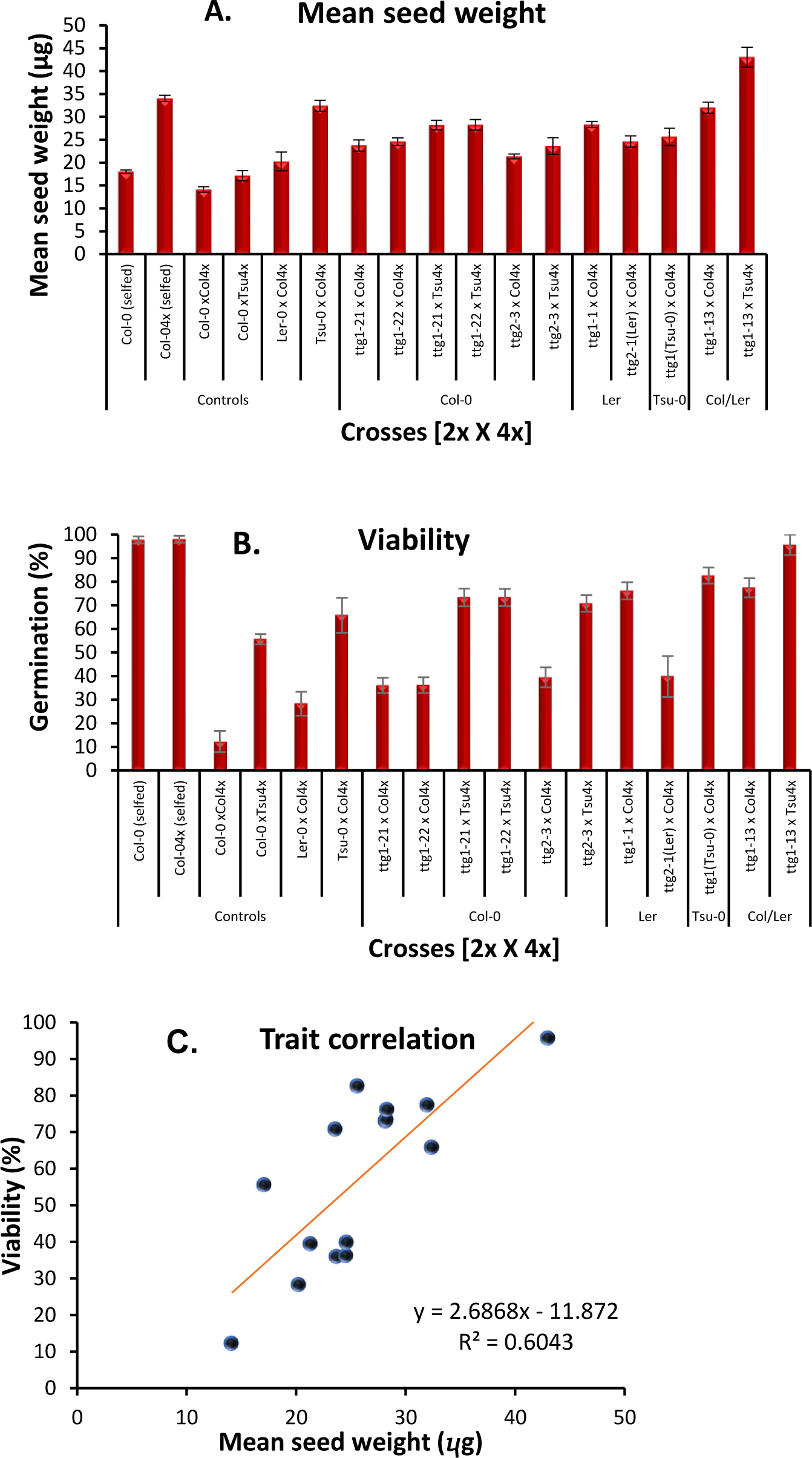
The maternal mutation of *TTG1* increases the viability of the triploid F_1_ seeds generated from 2x X 4x inter-crosses. A: Mean seed weight of the seeds derived from the interploidy crosses compared with different controls. Only the *ttg1* mutant in Tsu-0 has a significant reduction in mean seed weight. B: The percentage viability of the seeds derived from the interploidy crosses compared with the controls. The mother comes first in all crosses. C. A positive correlation between seed viability and mean seed weight in the interploidy crosses with *ttg1* mothers. The error bars are standard error of the mean.

All the maternal *ttg1* mutations significantly repressed seed lethality, with the seeds resulting from these inter-crosses exhibiting a mild or strong rescue (Figures 5 and 6). The weak rescuer group had a considerable percentage of shrivelled seeds (24µg mean seed weight and 40% viability) in response to Col4x pollen (Figures 5 and 6). These were obtained from the interploidy crosses featuring the *TTG1* and *TTG2* knockouts in the Col-0 background. *TTG1* knockouts in Ler (*ttg1-1)* and a Col-0/Ler F_3_ line (*ttg1-13*) resulted in increased seed survival (28µg mean seed weight and 78% viability) (Figures 5 and 6). Similarly, the *ttg1* mothers in a pure Col-0 background substantially bypassed the mild killing induced by Tsu4x pollen to produce a high percentage of viable seeds.

In contrast, the CRISPR-Cas9 knockout of the *TTG1* protein in Tsu-0 resulted in a significant reduction in mean seed weight compared to wildtype Tsu-0 (Figures 5 and 6). These results strongly indicate the influence of accession differences in the outcomes of the *TTG1* mutant mothers in interploidy crosses. Thus, the Col-0 wildtype *TTG1* allele is lethal for maternal rescue, whereas the Tsu-0 wildtype *TTG1* allele promotes maternal rescue against Co4x killing.

### *TTG1* synergistically interacts with *AXR1* to increase seed size in diploids

Triploid F_1_ seed abortion has been reported to be associated with increased auxin activity following fertilisation, and the downregulation of auxin biosynthesis and signalling genes improved seed survival in interploidy crosses (*36*). To investigate if the lowered auxin requirement for seed development could potentially increase seed size or the maternal rescue potential of *ttg1* and *ttg2* mutants in a Col-0 mother, two double mutants were made by crossing a homozygous *axr1-3* mutant with homozygous *ttg1* and *ttg2* mutants, respectively (Figure S2). The *auxin-resistant* line (*axr1-3*) was used as it has moderate morphological abnormalities when compared to other auxin mutants (*36, 37*). The F_1_ plants obtained from the *ttg1/2* and *axr1-3* crosses were selfed to obtain homozygous F_2_s that were *axr1*-3like, inherited a homozygous *axr1-3* mutation plus the corresponding *ttg1/3* mutation (Figure S2 and Figure S3). The *glabrous* phenotype was more pronounced in the *ttg1-22axr1-3*F_2_s with no trichomes, indicating a positive additive effect of *axr-1* and *ttg1* (Figure S2). In addition, the *ttg1-22axr1-3*F_2_s were more fertile and produced longer siliques to suppress the short siliques induced by the *axr1* gene, signifying a possible compensation strategy against the adverse pleiotropic effects of the *axr1* mutation (Figure S2).

Furthermore, these *ttg1-22axr1-3*F_2_s significantly produced bigger seeds similar to Col-0 tetraploids, and this increase is lacking in their respective parents and the *ttg2-3axr1-3*F_2_s (Figures 7A, B and C). The average seed area of a *ttg1* and *axr1* double mutant was approximately increased to 60% of the diploid wild-type (Figures 7A and C). The synergistic increase in seed size resulting from the double mutation in *ttg1* and *axr1* is remarkably distinguishable from the other diploids starting from 7 Days After Pollination (DAP), and this is consistent with an increased endosperm volume (Figures 7E and F). *TTG2* is significantly less expressed in different seed tissues when compared to *TTG1* and *AXR1,* suggesting distinct maternal roles in seed development (Figure 7D). However, the positive additive action of *ttg1* and a*xr1* in the other traits does not entirely abolish the submissiveness to Col4x-killing in a Col-0 mother (Figures 8A and B). Thus, these results strongly suggest that distinct auxin pathways mediate the *ttg1*/*ttg2* seed size regulation in diploids and triploids.

**Figure 7:**
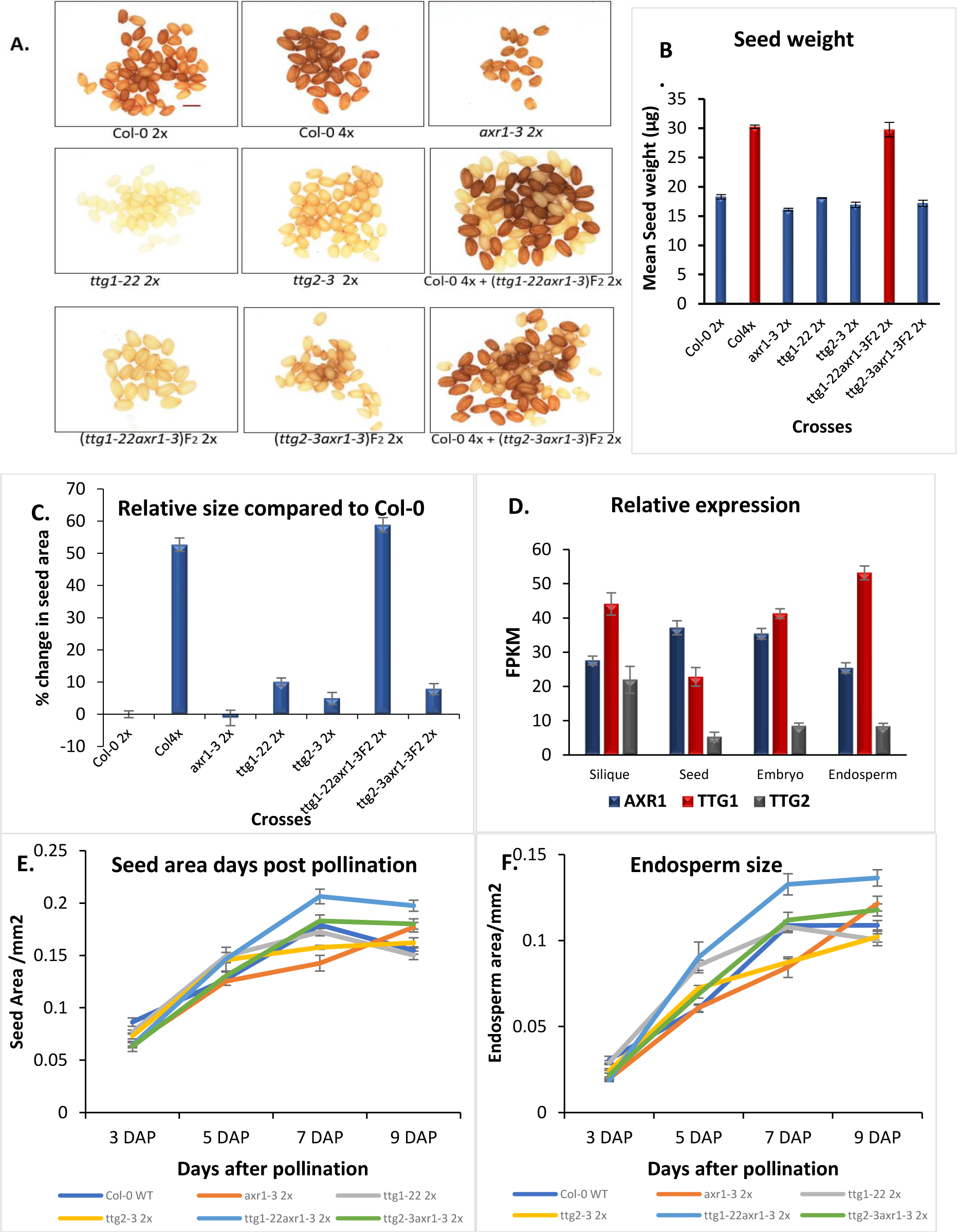
The *ttg1axr1* double mutants show a strong positive additive increase in seed size and weight that is lacking in *ttg2axr1* double mutants. A: Images of the mature diploid seeds obtained from *ttg1axr1*F_2_s compared with their parental controls and a Col-0 tetraploid. The diploid double mutants (*ttg1-22axr1-3* and *ttg2-3axr1-3,* respectively) were mixed with Col-0 tetraploids for seed size comparison. All the images were captured on the same scale to compare seed sizes. The scale bar is 0.5 mm in A. B: The mean seed weight of the *ttg1axr1* and *ttg2axr1* double mutants compared with their parental controls and a Col-0 tetraploid. C: The average seed weight of a typical *ttg1axr1* double mutant is similar to a Col-0 tetraploid. The bars on the positive axis indicate an increased seed size, while the bars on the negative axis indicate a decreased seed size. D: *TTG2* is relatively less expressed in seed tissues, unlike *TTG1* and *AXR1.* The data used for plotting were retrieved from 137 public transcriptome libraries in the Arabidopsis RNA-seq database (*38*). E and F: Comparing the seed size and endosperm volume of the mutants during the early stages of seed development. The error bars indicate the standard error of the mean.

**Figure 8:**
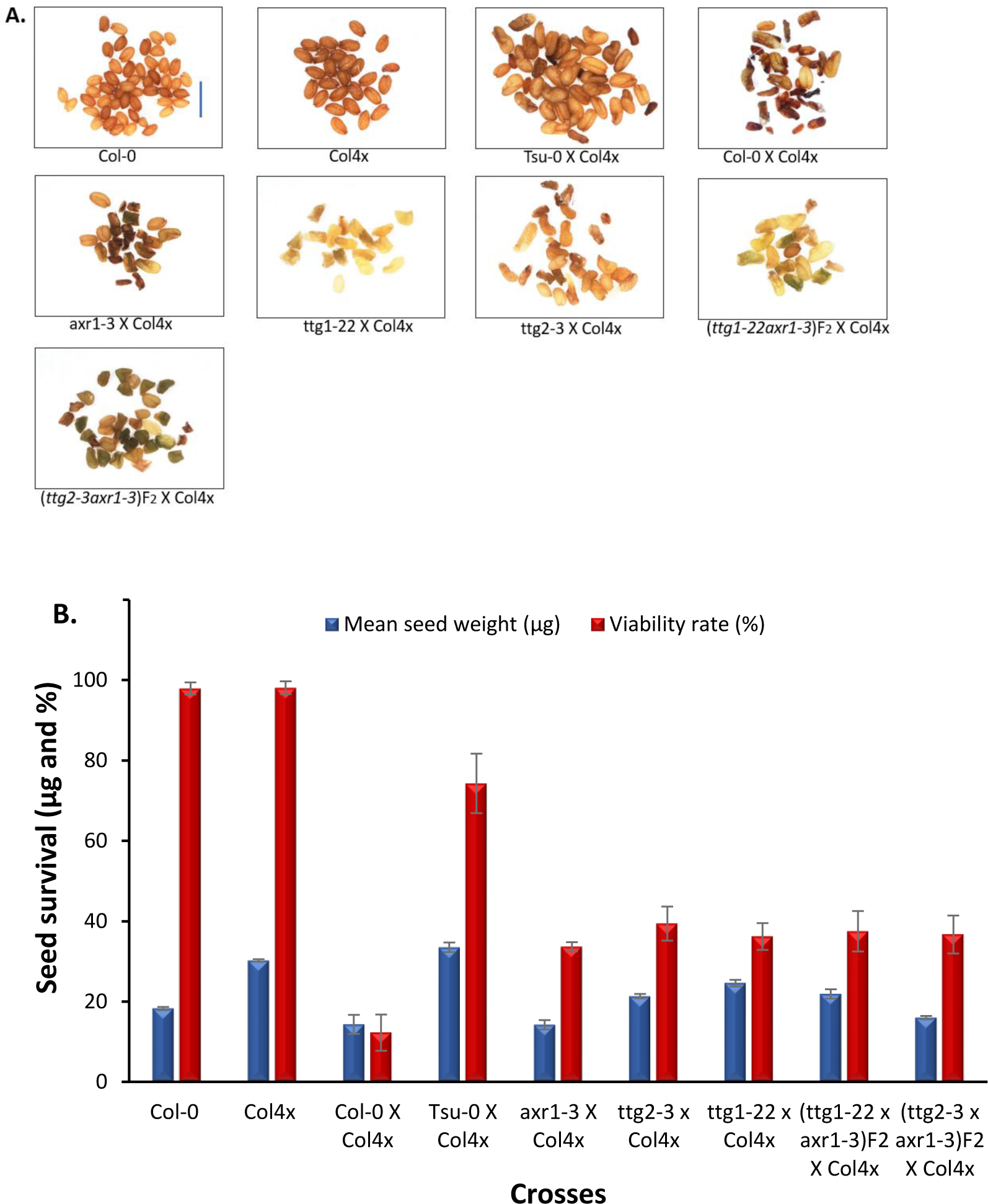
Maternal double mutations in *axr1* and *ttg1*/*ttg2* have a significant but weak additive effect on maternal rescue in a triploid block. A. The mean seed weight of the *axr1-3* double mutants in interploidy crosses B. The germination rate of the seeds obtained from interploidy crosses. The y-axis represents the level of seed survival in terms of mean seed weight (µg) and percentage viability rate. The x-axis represents the different interploidy crosses (2x X 4x). The error bars indicate the standard error of the mean. The scale bar is 1mm.

## DISCUSSION

The paternal and the maternal genomes play crucial roles in interploidy seed size and survival. Some genetic components such as *NRPD1*, *ADMETOS*, *SUVH7*, *SUVH9*, *AHL10*, *PEG2*, *PEG9, PICKLE RELATED2* (*PKR2)*, *PHERES1* (*PHE1*), *TTG2* and *TT4* have been validated to suppresses lethality in F_1_ triploids (*10, 18, 36, 39-44*). However, most of these genes are paternally expressed (PEGs) in the endosperm (*9, 39, 40, 45, 46*). An interplay of different mechanisms, such as different DNA methylation patterns, gene expression differences, allelic states, hormones and other maternal cues, underlie the mode of action of these genes. However, the broad natural variation in the interactions in these multiple loci, especially in different genetic backgrounds, remains unanswered.

This study showed that maternal cues such as *TTG1*, *TTG2* and auxin act in the different seed compartments to alleviate Col4x killing. Interestingly, *TTG1* and *TTG2* are involved in the transcriptional regulation of the FBP pathway, with *TTG1* acting as the conserved transcriptional head in all land plants (*22*). *TTG2* function strictly depends on an intact *TTG1* gene for its proper expression in the seed coat endothelium during early seed development (*28*). The maternal rescue action of these genes in the triploid block is accession-dependent. Some accessions like Bla-1 and Ler utilise *TTG2* on chromosome 2 as the primary maternal modifier of col-killing (*10*). Interestingly, we showed that other accessions like Tsu-0 map to the *TTG1*-locus on chromosome 5. However, mutations in all the maternal modifiers of the triploid block (*ttg1*, *ttg2* and *axr1*) do not completely alleviate Col-killing in Col-0. This strongly implies that an interplay between or among the MEGs and other loci acting in all the seed compartments is required to significantly improve maternal rescue in a typical Col-0 diploid. Genic interactions explain how these loci operate in different backgrounds, and these could range from additive to complementary, epistasis or even duplicate genes.

*TTG1* and *TTG2* are both responsive to auxin, a maternal signal to trigger endosperm cellularisation (*36, 47*). Auxin is produced in the endosperm and transported to the seed coat. Therefore, the downregulation of auxin-related gene expression in all the endosperm domains is crucial for endosperm cellularisation (*36, 48*). The lack of a high additive gene action of the *ttg1/ttg2-axr1-3* double mutants in abolishing 2x X 4x interploidy F_1_ lethality describes a common genetic link between the FBP pathway and auxin signalling in the triploid block. However, the data on the genetics of auxin-related seed size and the *TTG* double mutants in diploids indicates that *TTG1* and *TTG2* are neither equivalents nor interchangeable in diploid seed size control. While the *TTG1*/*TTG2* pathways may be connected in seed coat colour and trichome regulation, their roles in seed size in diploids have diverged in the FBP pathway. The divergence could have occurred via the auxin-responsive branch of the FBP featuring quercitrin and kaempferol (*49*). This suggests a complex, trait-dependent functional relationship between the *TTG1/TTG2* gene products and auxin during seed development.

*TTG1* and *TTG2* act in different auxin pathways to affect diploid seed size. The *TTG1* role in increasing diploid seed size is associated with auxin repression. The positive additive gene action recorded in the *ttg1axr1* diploid seeds may be linked to an increased accumulation of nutrients during early endosperm development or increased seed storage reserves at maturation. The latter corroborates a previous study demonstrating increased dry weight in *ttg1-1* embryos (Ler background) compared with their wildtype (*50*). However, the *TTG1-FUSCA3* mechanism is only a minor increase in seed size. Enolase2 has also been reported to play a role in fatty acid accumulation during seed maturation by interacting with flavonoids and auxin (*51*). Other mechanisms, such as delayed endosperm cellularisation, increased carbohydrate accumulation, prolonged seed maturation or alterations in some other signalling pathways controlling seed size in maternal tissues, could be responsible for the increased seed size in *ttg1axr1-3* double mutants.

In conclusion, *TTG1* is an essential determinant of seed size in both diploids and triploids. Since the *TTG1* and auxin pathways are conserved in all plants, we believe that reduced auxin signalling in a null *ttg1* background can provide a rapid strategy for increasing seed size in diploids.

## MATERIALS AND METHODS

### Plant materials and growth conditions

Prior to planting, seeds were stratified at 4°C as a suspension in 0.1% Phyto agar for 5 days except for the 2RV set of RILs that was stratified for longer (7days). All plants were grown in growth rooms (Sanyo) under a controlled humidity of 40%, temperature of 21°C, 16-hour day length and 8-hour night length. Following stratification, the seeds were planted on F2+S compost (Levingtons) that has been pre-treated with imidasect, a slow-release insecticide. Two distinct recombinant inbred line populations (RILs) were used for this study (*52*). The first core RIL population of 164 lines (3RV) were obtained from the French National Institute for Agricultural Research (INRA) and used the *A. thaliana* accessions, Col-0 (186AV) and Tsu-0 (91AV) as the original parent lines. The second RIL population (2RV) was also obtained from INRA and was developed from the Col-0 (186AV) and Bla-1 (76AV) accessions. All the Col4x seeds recruited as pollen parents were previously tetraploidised and verified by Wilkins (2021). A *ttg1* clustered regularly interspaced short palindromic repeats and CRISPR-associated protein 9 (CRISPR-CAS9) mutant line in Tsu-0 (N1564) genetic background previously generated in the plant lab, the University of Bath was also used. The rest of the lines used for this study are accessions and mutant lines obtained from the Nottingham Arabidopsis Stock Centre (Table S5).

### Controlled crosses and Seed phenotyping

All the plants used as mothers for each cross were emasculated and pollinated immediately with a high pollen dose to minimise the chances of cross-contamination. Stage 12 flower buds on the primary inflorescence of the seed parent were emasculated, and the remaining flowers and buds around the emasculated buds removed. Mature pollen from the appropriate pollen parent was gently dabbed on the surface of the stigma of the emasculated buds. Both emasculation and pollination were carried out using a pair of sharp tweezers with the aid of a x3.5 magnification optivisor. Mature brown siliques were removed from each seed plant around 3-4 weeks after each pollination event. Each silique was gently opened, all the seeds were collected separately and viewed using a Nikon SMZ1500 Microscope and the NIS-Elements AR software (Nikon, UK). The images were captured using a Nikon Digital Sight DS-U1 camera. The weight of the seeds was measured in µg using the Mettler UMT2 microbalance (Mettler-Toledo, Leicester, UK). Plump and shrivelled seeds were categorised in each photographic image using the image J counter tool (*53*). Seed size analysis was done with the image J particle analysis tool.

### Seed Viability

The seed germination protocol used was adapted from (Podar, 2013). All seeds were sterilised with 1% chlorine gas in a closed chamber for 1 hour. The resultant were evenly placed on agar plates containing Murashige Skoog growth medium with MES buffer (Duchefa Biochemie/Melford), 0.8% Phyto Agar (Duchefa Biochemie/Melford) and 0.1% sucrose (Acros) (final pH of 5.6-5.7). The plate with four replications sseeds were cold stratified at 4oC for two days on agar and later placed in growth rooms (Sanyo) under a controlled humidity of 40%, temperature of 21°C, 16-hour day length and 8-hour night length to allow for radicle emergence. A hundred Col-0 tetraploid seeds in four separate plates were used as the control plates to normalise the viability data.

### QTL analysis

The overall mean of the seed weight measurements resulting from each diploid RIL crossed with a Col4x father in four replicates was used as the phenotype data. Genotype data for all the RILs was downloaded from the INRA webpage. WinQTL cartographer (version 2.5) was used to calculate log-likelihood (LOD) ratio statistics using composite interval mapping. The mapping parameters were set to a statistical threshold with a significance level of p<0.05 using 1,000 permutations for each trait.

### Graphical analysis

All graphs were plotted using the R software, SPSS (version 25), GraphPad prism 8, Microsoft Excel 365 and ChiPlot (https://www.chiplot.online/#Line-plot). Where applicable, the standard error of the mean or the standard deviation were calculated using the R software. **Indel marker development and genotyping**

The Integrative Genomics Viewer (IGV) was used to visualise indels and SNPs. TAIR 10 (default genome for *A. thaliana* on IGV) was used for the Col-0 accession while a BAM file containing the Tsu-0 illumina reads (mapped to the Col-0 reference genome) was retrieved from the European Nucleotide Archive. The correct identifier for each bam file and bai file was obtained from the 1001 Genomes website (http://1001genomes.org).

Leaf samples were collected from three to four weeks old seedlings, with the sample material being selected from younger, freshly expanded leaves. All the leaf samples were stored at - 20°C prior to DNA isolation. A 20 µl reaction volume was set up for each PCR consisting of 0.5 µl of each DNA supernatant, 1.0 µl of each primer, 7.5 µl nuclease-free water and 10.0 µl of the Phire Plant PCR Mastermix. A no-template control was included in all reactions to check for contamination. All PCR reactions were run in a GSTORM thermocycler using the following programme: initial denaturation at 98°C for 5 minutes, final denaturation at 98°C for 5 seconds, annealing temperature (specific to each primer) for 5s, extension at 72°C for 20s (time per Kb of each expected PCR product) and a final extension at 72°C for 1 minute. The total number of cycles for denaturation, annealing and extension was 40. Following PCR, products were loaded on 2.5 % agarose gels containing Ethidium bromide (1 µl of Ethidium bromide to 35 µl of each gel). Each gel was run for 35minutes with 100V and 2.5A. Polymorphisms in the PCR products were visualised and captured after electrophoresis using a UV transilluminator.

### Candidate gene analysis

To define the genomic interval containing the candidate genes, the genotype data were first sorted with the phenotype data (from low to high), and the approximate interval was determined by the chromosomal position (in Mb) of the two markers at the boundary of the QTL. The total number of genes within the narrowed QTL intervals were retrieved from the *A. thaliana* EPD Viewer Hub Assembly with UCSC table browser data retrieval tool (*54, 55*). Different gene models, gene alias, all gene ontology terms, functional descriptions, curator summary and publication links for all the retrieved candidate genes were downloaded from the TAIR 10 gene ontology (GO) and plant ontology (PO) annotations available on the TAIR website (https://www.arabidopsis.org/download/index-auto.jsp?dir=%2Fdownload_files%2FGO_and_PO_Annotations%2FGene_Ontology_Annotations). The first and the last genes to appear on the gene list from the UCSC genome browser served as the coordinates used to parse the data downloaded from TAIR.

All the GO terms for all gene models for the gene list obtained from the UCSC genome browser were analysed on agriGO (version 1.2), a GO analysis toolkit(*56*). A singular enrichment analysis (SEA) was performed using this gene list as the query list for the *A. thaliana* TAIR 10 genome locus (TAIR 10, 2017). Fisher exact tests and Hochberg multi-test adjustment method were applied as implemented in the R-function fisher.exact and R-function p. adjust respectively at a significant level of 0.01. A FDR (false discovery rate) of 0.01 was also applied to define the level of stringency of the Hochberg multiple testing correction applied to the Fisher exact test results. Significant associations between the total number of traits curated from the gene list and the biological process (BP) associated was derived based on the overrepresentation of BP terms within the input list using a p-value of 0.01 and the FDR. The significant GO lists were summarised with REVIGO and visualised with Chiplot.

### CRISPR-Cas9 gene editing

The Tsu-0 accession (NI564) obtained from NASC is the original genetic background of the CRISPR-Cas9 *ttg1* mutant line. CHOPCHOP Harvard (version 3) was used to design appropriate two 19 nucleotide guide RNAs close to the promoter region of *A. thaliana TTG1* (AT524520.1, isoform 1). A *pCBC-DT1DT2* plasmid obtained from Addgene was used as the template for the assembly of two gRNA expression cassettes using a four-primer mixture in a high-fidelity PCR with Phusion High-Fidelity DNA Polymerase (Table S1). The constructs were assembled into the *pHEE401E* Cas9 vector via golden gate cloning. The plants were transformed via Agrobacterium-mediated transformation by floral dipping and the primary transformants verified with BASTA treatment. Following stratification, the seeds were planted on F2+S compost (Levingtons) that has been pre-treated with imidasect, a slow-release insecticide. All plants were grown in growth rooms (Sanyo) under a controlled humidity of 40%, temperature of 21°C, 16-hour day length and 8-hour night length.

Each seed batch obtained from the Agrobacterium transformed plants were directly sown on soil in hundreds by broadcasting and cold stratified in the dark for five days with wildtype and BASTA resistant controls. Ten days after cold stratification, all the seedlings were treated with 50 mg/L of BASTA (Glufosinate-ammonium, SIGMA). The BASTA treatment was repeated on the eleventh and twelfth days. All the successfully transformed plants with green leaves were nurtured till maturity, and the dry seeds were harvested per individual. Young leaves were collected from all the surviving seedlings for DNA isolation. A set of primers flanking the vicinity of the guide RNAs designed were generated using CHOPCHOP Harvard (Table S6). The *TTG1* PCR fragments amplified from the genomic DNA of these leaf samples were sequenced identify chimaeras in the T_0_ generation. The chimaeras identified were individually selfed to produce a T_1_ population. All the selfed plants displaying obvious *TTG1* mutations on the leaves, stems and seeds were carefully identified within the T_2_ generation. The accuracy and position of CRISPR-cas9 deletions within the *TTG1* gene of each successful transformation was verified by sequencing. The resultant ttg1(Tsu-0) mutant has been donated to NASC with the accession number: N2111669 and ABRC stock number: CS2111669.

