## Supplemental information for "Busted: maternal modifiers of the triploid block involved in seed size control"

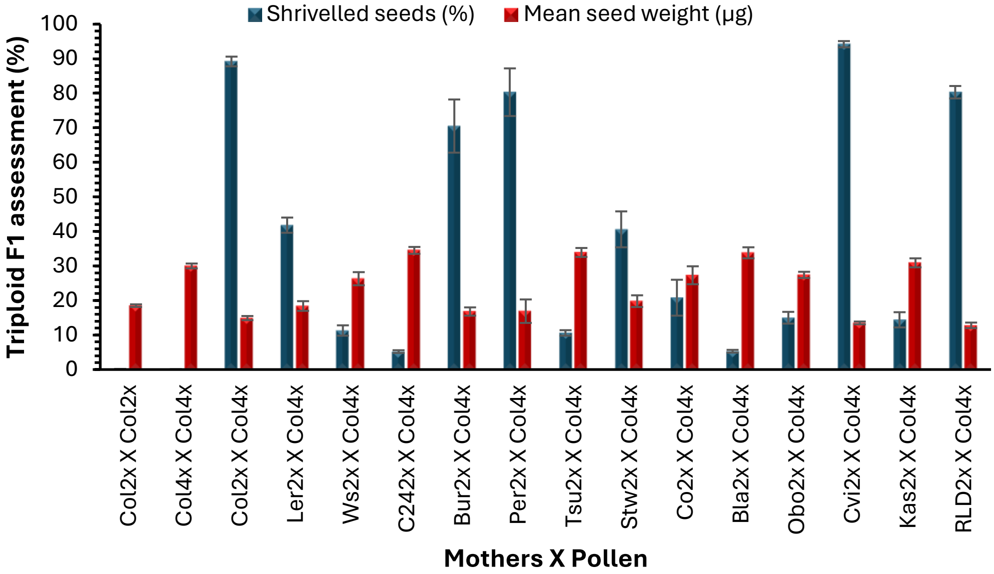


**Figure S1: Maternal rescue against Col4x pollen in Arabidopsis is accession-dependent.** Some accessions like Col, Cvi and RLD are poor maternal rescuers in response to Col4x pollen resulting in a high percentage of shrivelled seeds. Other accessions like Tsu-0, C24, Bla-1, Kas, Per and Ws maternally resist the killing activity of Col4x to produce viable offspring, with high mean seed weight. The error bars are standard error of the mean. (Adapted from Bolbol, 2010).

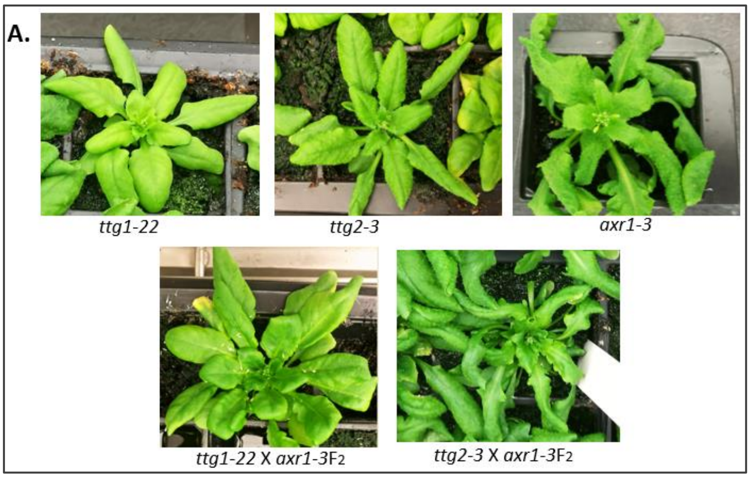


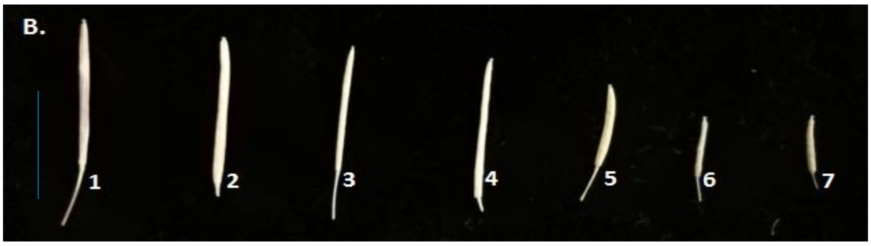


**Figure S2: The phenotypes of *axr-1* and *ttg1/ttg2* double mutants.** A. Images of the plants at the bolting stage (Note that the flowering times are different). B. Mature siliques with dry seeds of 1(Col-0), 2(Col-04x), 3(*ttg2-3*), 4 (*ttg1-22*), 5(*ttg1-22* X *axr1-3*F2), 6(*ttg2-3* X *axr1-3*$F_2$), and 7(*axr1-3*). The Scale bar is 10mm in B.

**A.**

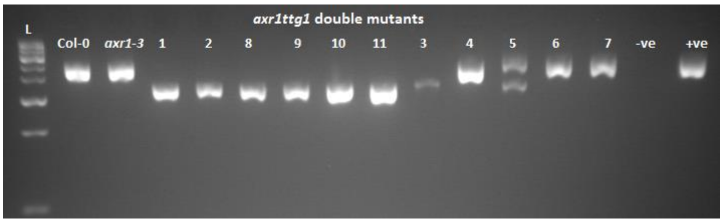


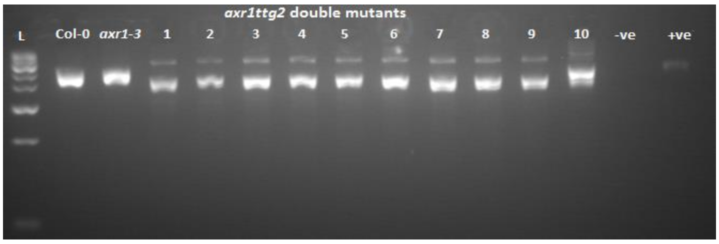


**B.**

```
Col-0               TCGAATACAGAGCTCGTAAAGAAAAAGAAGGCAAAAGCTTCAAAAACCACGTCATTCAGG      214
axr1-3              TCGAATACAGAGCTCGTAAAGAAAAAGAAGGCAAAAGCTTCAAAAACCACGTCATTCAGG      417
axr1-3ttg1-22       TCGAATACAGAGCTCGTAAAGAAAAAGAAGGCAAAAGCTTCAAAAACCACGTCATTCAGG      418
axr1-3ttg2-3        TCGAATACAGAGCTCGTAAAGAAAAAGAAGGCAAAAGCTTCAAAAACCACGTCATTCAGG      420
                    ************************************************************

Col-0               TCCCCTCTTTAAACCCTCTCTCTCTAATCAGAATCACCAAAACATAATTATCTTTCACTA      274
axr1-3              TCCCCTCTTTAAACCCTCTCTCTCTAATCAGAATCACTAAATCATAATTATCTTTCACTA      477
axr1-3ttg1-22       TCCCCTCTTTAAACCCTCTCTCTCTAATCAGAATCACTAAATCATAATTATCTTTCACTA      478
axr1-3ttg2-3        TCCCCTCTTTAAACCCTCTCTCTCTAATCAGAATCACTAAATCATAATTATCTTTCACTA      480
                    ************************************* *** ******************

Col-0               ATCTAACACTTTTAAAAATCTCAACAGTGGTTTGAAGTACTTGATGGATTACTTGGTTCA      334
axr1-3              ATCTAACACTTTTAAAAATCTCAACAGTGGTTTGAAGTACTTGATGGATTACTTGGTTCA      537
axr1-3ttg1-22       ATCTAACACTTTTAAAAATCTCAACAGTGGTTTGAAGTACTTGATGGATTACTTGGTTCA      538
axr1-3ttg2-3        ATCTAACACTTTTAAAAATCTCAACAGTGGTTTGAAGTACTTGATGGATTACTTGGTTCA      540
                    ************************************************************
```

**C.** Primers

| Primer | Primer sequence | Phire Tm ($^{0}$C) |
|---|---|---|
| ***axr1-3-F*** | **GTCGGAAGGAGTAGAAGCGA** | 62.6 |
| ***axr1-3*-R** | **TTCAACGGTCTTGTGTGTCT** | |
| ***ttg1-22* LP-F** | **TCGATTGGAACGATGTAGAGC** | 61.0 |
| ***ttg1*-22 RP-R** | **TGGGTAAAAATTAGAACCTGACG** | |
| **GABI-F** | **ATATTGACCATCATACTCATTGC** | |
| ***ttg2-3*F LP-F** | **TAAAACCAAACGACACCGTTC** | 61.3 |
| ***ttg2-3*R RP-R** | **TCCAAGTTTGTTGACGATTCC** | |
| **SALK-F** | **ATTTTGCCGATTTCGGAAC** | |

**Figure S3: The *axr1* and *ttg1/ttg2* $F_2$s were confirmed to be double mutants**. A: Confirmation of the *ttg1/ttg2* alleles by PCR. The numbered individuals in both gels are the $F_2$s and are ranked from the left panel to the right panel by the strength of the phenotypes. B: Confirmation of the *axr1-3* allele by sequencing. The area with no star indicates the nucleotide substitution characterising the *axr1-3* mutation. C: A list of the primers used for confirming the mutations. The *ttg1* primers are GABI primers, while the *ttg2* primers are SALK primers. *axr1-3*F and *axr1-3*R were designed in this study and used to amplify a 4.5 Kb fragment of the *axr1-3* gene. *axr1-3*F was used as the sequencing primer to verify the EMS mutation.

### Table S1: The foreground indel markers designed within the Tsu-0 QTL interval

| Primer name | Sequence | Product size in Colombia (bp) | Product size in Tsu-0 (bp) | Tm (˚C) |
|---|---|---|---|---|
| 4.03 | F: ACAAGTGACCGATCCAATTTGT<br>R: TGTGCCGCATGTAACATAAACT | 494 | 194 | 67.4 |
| 4.40 | F: CCATGCCCTTTGATCATCTGA<br>R: ATTGAGAGAGTCCTGCCCAC | 354 | 250 | 63.7 |
| 4.94 | F: ATTGGGCCCGTTTTGAGATG<br>R: TCGTCAGAGTGTTGTTCCGA | 794 | 600 | 65.1 |
| 5.48 | F: GGGCCTTAACCGGAGATTATT<br>R: GGTTCTTGTTGGGTCGCAA | 272 | 122 | 64.2 |
| 5.96 | F: ATCCAGTCCACATCAACCCT<br>R: CTCCACCACAGCAGCCTC | 5,050 | 700 | 63.7 |
| 6.50 | F: TCGTGTAAGCTGTTTCTTCGA<br>R: ACTCGCTCATCAAATCACCTC | 226 | 100 | 65.9 |
| 7.20 | F: TGTGTAGGAGGACGATGATGA<br>R: AGCTGAAATCCACAATGCGA | 174 | 80 | 63.6 |
| 7.75 | F: ACACAATCCTCCTCTCCACT<br>R: TCCGACTGTGATTGGACCC | 956 | 550 | 60.7 |
| 8.20 | F: AACTTCTCCACACGGTAGGG<br>R: GCCACGGAAGAAGAAAGACC | 695 | 395 | 63.3 |
| 8.32 | F: CCAATGGTTGAGTACGCGAT<br>R: GCACTCTGAGGGGATTAGGG | 1567 | 680 | 63.4 |
| 8.58 | F: CCAAGACCAGGGGAGTGATT<br>R: CACTCGTACTGTGCTCTTGT | 1786 | 650 | 62.6 |

Tm, annealing temperature; bp, base pairs; F, forward; R, Reverse. The markers are arranged from the beginning to the QTL peak.

**Table S2: A list of all the significant GO terms enriched in the candidate genes within the narrowed QTL interval.**

| GO CATEGORY | GO Term | Description | Log10 p-value | Uniqueness | Dispensability | FDR |
|---|---|---|---|---|---|---|
| BP | GO:0045927 | positive regulation of growth | -10.0969 | 0.836 | 0 | 8.00E-11 |
| | GO:0035315 | hair cell differentiation | -10.0362 | 0.629 | 0 | 9.20E-11 |
| | GO:0010026 | trichome differentiation | -10.0362 | 0.59 | 0.52 | 9.20E-11 |
| | GO:0045793 | positive regulation of cell size | -9.7696 | 0.747 | 0.13 | 1.70E-10 |
| | GO:0009965 | leaf morphogenesis | -8.1871 | 0.617 | 0.348 | 6.50E-09 |
| | GO:0007049 | cell cycle | -4.3665 | 0.784 | 0.223 | 4.30E-05 |
| | GO:0051276 | chromosome organisation | -4 | 0.828 | 0.261 | 0.0001 |
| | GO:0008361 | regulation of cell size | -3.5376 | 0.713 | 0.905 | 0.00029 |
| | GO:0090066 | regulation of anatomical structure size | -3.5376 | 0.882 | 0.36 | 0.00029 |
| | GO:0006259 | DNA metabolic process | -3.5086 | 0.892 | 0.259 | 0.00031 |
| | GO:0040007 | Growth | -3.1024 | 0.97 | 0 | 0.00079 |
| | GO:0032502 | developmental process | -2.9208 | 0.973 | 0 | 0.0012 |
| | GO:0032501 | multicellular organismal process | -2.4437 | 0.973 | 0 | 0.0036 |
| | GO:0009607 | response to biotic stimulus | -2.4318 | 0.917 | 0.335 | 0.0037 |
| | GO:0008104 | protein localisation | -2.0223 | 0.953 | 0 | 0.0095 |
| | GO:0051704 | multi-organism process | -2.0088 | 0.97 | 0 | 0.0036 |
| CC | GO:0009330 | DNA topoisomerase complex (ATP-hydrolysing) | -10.4318 | 0.44 | 0 | 3.70E-11 |
| | GO:0032991 | macromolecular complex | -1.7212 | 0.62 | 0 | 0.0068 |
| MF | GO:0003690 | double-stranded DNA binding | -6.8539 | 0.75 | 0 | 1.40E-07 |

BP-Biological process, CC-Cellular component, and MF-molecular function. Most of the non-significant GO entries have been filtered out with p-values (>0.05) and FDR and do not appear in this table and further analyses. Log10 p-value is derived from the logarithm value of each GO entry.

Definition of terms

1. Dispensability: The semantic similarity threshold at which the term was removed from the list and assigned to a cluster. Cluster representatives always have dispensability less than the user-specified 'allowed similarity' cutoff.
2. Uniqueness: Measures whether the term is an outlier when compared semantically to the whole list, calculated as 1-(average semantic similarity of a term to all other terms). More unique terms tend to be less dispensable.

**Table S3: The final list of genes from the filtered gene set.**

| Gene | Gene alias | GO terms | Variants | Key traits | Expressed in seeds? |
|---|---|---|---|---|---|
| **AT5G24630** | *BIN4 (BRASSINOSTEROID-INSENSITIVE4)* | GO:0051276-Chromosome organisation<br>GO:0009330-DNA topoisomerase complex (ATP-hydrolysing)<br>**GO:0010090-Trichome morphogenesis**<br>**GO:0030307-Positive regulation of cell growth**<br>GO:0042023-DNA endoreduplication<br>GO:0048364-Root development<br>GO:0003690-Double-stranded DNA binding | AT5G24630.1<br>AT5G24630.2<br>AT5G24630.3<br>AT5G24630.4<br>AT5G24630.5<br>AT5G24630.6 | Dwarfism, reduced cell size in leaves, roots and hypocotyls, root hair growth, and trichome development. | Yes |
| **AT5G24520** | *TTG1 (TRANSPARENT TESTA GLABRA 1)* | **GO:0009965-leaf morphogenesis**<br>**GO:0009957-epidermal cell fate specification**<br>GO:0035315: Hair cell differentiation<br>GO:0000166-nucleotide binding<br>GO:0005515-protein binding<br>GO:0003677-DNA binding<br>**GO:0010026-trichome differentiation**<br>GO:0032880-regulation of protein localisation | AT5G24520.1<br>AT5G24520.2<br>AT5G24520.3 | Trichome differentiation, root hair growth, epidermal cell fate specification, flavonoid biosynthesis, mucilage deposition, anthocynanin content and proanthocyanidin deposition. | Yes |
| **AT5G24330** | *ATXR6 (ARABIDOPSIS TRITHORAX-RELATED PROTEIN 6)* | GO:0051726-regulation of cell cycle<br>GO:0005634-nucleus<br>GO:0006355-regulation of transcription, DNA-dependent<br>GO:0009901-anther dehiscence<br>GO:0005515-protein binding<br>GO:0003677-DNA binding | AT5G24330.1 | Chromatin structure and gene silencing, cell cycle regulation, anther dehiscence and petal differentiation | Yes |

The GO terms in bold are often related and control the same traits such as trichome differentiation and leaf morphogenesis.

**Table S4: Chi-square tests linking c5_8.32 and c5_8.58 to maternal rescue**

**A.**

| | Markers | AA | AB | BB | χ2 *p*-value (AA=BB) | $BC_1F_2$ fits into the expected ratio (AA=BB)? |
|---|---|---|---|---|---|---|
| 4.03Mb | c5_4.03 | 7 | 4 | 5 | 0.264 | Accept |
| 4.49 Mb | c5_4.4 | 4 | 11 | 1 | 0.004 | Reject |
| 4.94 Mb | c5_4.94 | 16 | 0 | 0 | 0.0001 | Reject |
| 5.48 Mb | c5_5.96 | 2 | 8 | 6 | 0.025 | Reject |
| 6.50 Mb | c5_6.50 | 14 | 0 | 2 | 0.006 | Reject |
| 7.20 Mb | c5_7.20 | 16 | 0 | 0 | 0.0001 | Reject |
| 7.75 Mb | c5_7.75 | 16 | 0 | 0 | 0.0001 | Reject |
| 8.20 Mb | c5_8.20 | 16 | 0 | 0 | 0.0001 | Reject |
| 8.58 Mb | c5_8.58 | 8 | 5 | 3 | 0.077 | Accept |

**B.**

| | | Observed | | Expected | | |
|---|---|---|---|---|---|---|
| | Locus | AA | BB | (AA=BB) | χ2 *p-value* (df=1) | $BC_1F_2$ expected ratio (AA=BB)? |
| **Submissives** | | | | | | |
| | C5_4.03 | 2 | 3 | 2.5 | 0.554 | Accept |
| | C5_8.58 | 7 | 1 | 4 | 0.034 | Reject |
| **Rescuers** | | | | | | |
| | C5_4.03 | 6 | 2 | 4 | 0.157 | Accept |
| | C5_8.58 | 7 | 1 | 4 | 0.034 | Reject |

**C.**

| | Markers | AA | AB | BB | χ2 p-value (AA=BB) | $BC_1F_3$ fits into the expected ratio (AA=BB)? |
|---|---|---|---|---|---|---|
| 4.03Mb | c5_4.03 | 19 | 7 | 6 | 0.01 | Reject |
| 4.49Mb | c5_4.49 | 4 | 10 | 17 | 0.01 | Reject |
| 4.94 Mb | c5_4.94 | 27 | 1 | 4 | 0.0001 | Reject |
| 5.48 Mb | c5_5.48 | 5 | 7 | 20 | 0.01 | Reject |
| 6.50 Mb | c5_6.50 | 31 | 0 | 1 | 0.0001 | Reject |
| 7.20 Mb | c5_7.20 | 31 | 0 | 1 | 0.0001 | Reject |
| 7.75 Mb | c5_7.75 | 31 | 0 | 1 | 0.0001 | Reject |
| 8.20 Mb | c5_8.2 | 31 | 0 | 1 | 0.0001 | Reject |
| 8.32 Mb | c5_8.32 | 15 | 4 | 13 | 0.4292 | Accept |
| 8.58 Mb | c5_8.58 | 13 | 8 | 11 | 0.145 | Accept |

**D.**

| | Locus | AA | BB | Expected(AA=BB) | χ2 *p-value* (df=1) | $BC_1F_3$ expected ratio (AA=BB)? |
|---|---|---|---|---|---|---|
| **Submissives** | | | | | | |
| | C5_8.58 | 10 | 2 | 6 | 0.021 | Reject |
| | C5_8.32 | 14 | 0 | 7 | 0.0001 | Reject |
| **Rescuers** | | | | | | |
| | C5_8.58 | 2 | 8 | 5 | 0.058 | Accept |
| | C5_8.32 | 0 | 12 | 6 | 0.001 | Reject |

Tables A and B are for $BC_1F_2$ progenies while tables C and D are for $BC_1F_3$ progenies. AA: Col-0 alleles; AB: Heterozygous; and BB: Tsu-0 alleles. Expected $F_2$ segregation ratio (1AA:2AB:1BB).

**Table S5: A list of the plant materials used in this study.**

| Type | Allele | Accession number | Accession | Mutation | Mutagenesis | Phenotypes |
|---|---|---|---|---|---|---|
| Wildtype | *NA* | N1564 | Tsu-0 | *NA* | *NA* | *NA* |
| | *NA* | N1092 | Col-0 | *NA* | *NA* | *NA* |
| | *NA* | NW20 | Ler-0 | *NA* | *NA* | *NA* |
| | *NA* | N970 | Bla-1 | *NA* | *NA* | *NA* |
| Mutants | *ttg1-1* | N89 | Ler | Q317stop | EMS | *tt, g* |
| | *ttg1-21* | N2105595 | Col-0 | Exon1 (N) | TDNA | *tt, g* |
| | *ttg1-22* | N2105596 | Col-0 | Exon1 (C) | TDNA | *tt, g* |
| | *ttg1-13* | CS67772 | Col-0/Ler $F_3$ | Whole deletion | Fast neutrons | *tt, g* |
| | *ttg2-1* | N277 | Ler | Exon1 (N) | TDNA | *tt, g* |
| | *ttg1-18* | N372 | En-1 | S310stop | Kranz collect | *NA* |
| | *ttg1-19* | N406 | Est-1 | W183stop | Kranz collect | *NA* |
| | *ttg1-15* | N300 | An-1 | S310stop | Kranz collect | *NA* |
| | *ttg1-Tsu-0* | N2111669 | Tsu-0 | Deletions | CRISPR-cas9 | *tt, g* |
| | axr1-3 | N3075 | Col-0 | - | EMS | |

Note that *ttg1-21* and *ttg1-22* are *ttg1* homozygous lines from the complete set of *tt*-isogenic collection lines (N2105571) from NASC. The other *ttg1* mutants were derived from (Larkin et al., 1999) and obtained from NASC. N277 is a *ttg2* mutant line. L*er*: Landsberg *erecta,* Col-0: Columbia-0, En-1: Enkheim-1, Est-1: Estland-1, An-1: Antwerp-1, EMS: Ethyl methanesulfonate (mutagenesis method), Q: Glutamine, S: Serine, W: Tryptophan, N: N-Terminus, C: C-Terminus, T-DNA: transfer-DNA, NA: not available as the lines failed to germinate, *tt*: *transparent glabra* and *g*: *glabra*.

**Table S6: Primer pairs for genotyping the CRISPR *ttg1*(Tsu-0) mutant**

| Primer name | Phire Tm | Sequence |
|---|---|---|
| 14 AND 1F | 60.8 | AAAATCCGACTGACACTGACCT |
| 14 AND 1R | | ATTGAATCGGAATCGAAAGAGA |
